# Optimal Binning of Peri-Event Time Histograms Using Akaike Information Criterion

**DOI:** 10.1101/2020.02.06.937367

**Authors:** Ali Ghazizadeh, Frederic Ambroggi

## Abstract

Peri-event time histograms (PETH) are widely used to study correlations between experimental events and neuronal firing. The accuracy of firing rate estimate using a PETH depends on the choice of binsize. We show that the optimal binsize for a PETH depends on factors such as the number of trials and the temporal dynamics of the firing rate. These factors argue against the use of a one-size-fits-all binsize when making PETHs for an inhomogeneous population of neurons. Here we propose a binsize selection method by adapting the Akaike Information Criterion (AIC). Simulations show that optimal binsizes estimated by AIC closely match the optimal binsizes using mean squared error (MSE). Furthermore, using real data, we find that optimal binning improves detection of responses and their dynamics. Together our analysis strongly supports optimal binning of PETHs and proposes a computationally efficient method for this optimization based on AIC approach to model selection.

## Introduction

The rate code theory of the nervous system proposes that neurons encode information about external stimuli via changes in their firing rate. To estimate a neuron’s response to an event, its spike trains from multiple occurrences of that event are aligned and spikes that occur around that event are counted in short time bins. These count histograms are then normalized by the number of trials and by the bin durations. This results in a peri-event time histogram or PETH which estimates the firing rate in Hz. One of the parameters that can affect the accuracy of a PETH in estimating the firing rate is the choice of binsize. However the choice of binsize is often left to the experimenter’s subjective assessment. Furthermore, in spite of the fact that noise levels and firing dynamics in different neurons warrants using a different binsize for each PETH, it is a common practice to use the same binsize for all the neurons recorded in a task. Importantly, this one-size-fits-all procedure ignores the inherent differences in the temporal resolution of PETHs, which can negatively affect the conclusions drawn from the population analysis of the neuronal activity by increasing response detection error and errors in estimating the response dynamics such as response onset and duration.

Here, we propose that the search for the optimal binsize can be viewed as a special case of the model selection problem where the number of bins used to estimate the firing represents the number of model parameters. Obviously, in a scheme with constant binsize over time the number of bins will also reflect the binsize. As for all model selection problems, having a large number of parameters (or small binsizes) results in over-fitting the data, while not having enough parameters (or large binsizes) fails to capture the variance of the data. Here we develop and propose a method based on Akaike information criterion (AIC) to determine the optimal binsize for each PETH (Akaike 1971). Our simulation results indicate that in most cases this method provides an accurate estimate of the optimal binsize which minimizes the mean squared error (MSE) between the firing rate estimate and the true firing rate.

## Methods

### Model Selection Approach to Binning

Let us assume that we have multiple trials of a neuron’s firing with the same underlying rate *λ*(*t*) in each trial. Our goal is to find 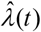 which minimizes the mean squared error or MSE defined as following:

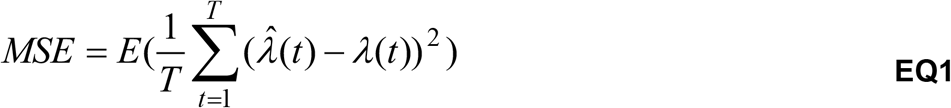

Where *E* is with respect to the distribution of 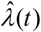 and *T* represents the trial duration in samples where time is considered to be a discrete variable which depends on the sampling rate. Minimizing MSE with respect to 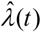 should result in a minimum variance estimate of the true firing rate during a trial.

The difficulty of minimizing MSE arises from the fact that in dealing with actual data *λ*(*t*) in not known. Therefore, one needs to find an approximate way to minimize the MSE.

Here our model space includes the PETHs with all possible binsizes used to estimate *λ*(*t*). We have:

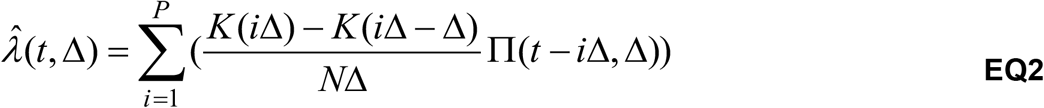

Where Δ refers to the binsize and 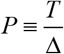 is the total number of bins for trial duration *T*. We define *K*(*t*) as the total number of spikes in all the *N* trials from the beginning of the trials up-to time *t* and define Π(*t*) as following:

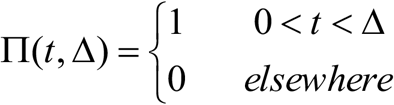

Using the PETH formulation presented in **EQ2**, we observe that the total number parameters that are used to construct the PETH is equal to the number of bins. Therefore the problem of choosing the best binsize which minimizes MSE can be viewed as a model selection problem with different number of parameters being the number of bins.

Among the various model selection methods in statistics AIC is one that is probably the most popular and at the same time the most effective for a vast range of model selection problems (De Leeuw 2007). Therefore we have adapted this method for choosing the optimal binsize. In deriving the AIC formulation for our current problem we assumed that the estimation errors are close to normally distributed in which case we have:

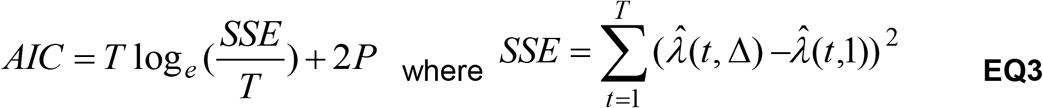

Here 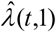 represents the firing rate estimate with the smallest binsize equal to 1 sample unit. 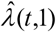 is taken to be the model free raw data that is being estimated by 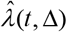. SSE is minimized when Δ is goes to one in which case the number of parameters *P* is as large as the number of samples or *T*. However since AIC includes a term that penalizes large parameter numbers in most cases AIC is only minimized at intermediate binsizes that result in lower number of parameters compared to *T*. This is because very small binsizes can result in over fitting the data and usually are not optimal.

In the simulation results we will demonstrate that in fact the AIC formulation presented in **EQ3** behave similarly to MSE as a function of binsize over a wide range of experimental conditions including number of trials and firing rate changes. The AIC cost used here is filtered by a 10 point moving average to yield a smooth trend as a function of binsize and to reduce the jitter in finding the optimal binsize.

### Shimazaki-Shinomoto cost function

A cost function based on the MSE for firing rates that follow Poisson distribution can be explicitly derived (Shimazaki and Shinomoto 2007). This cost function adapted to our notation is:

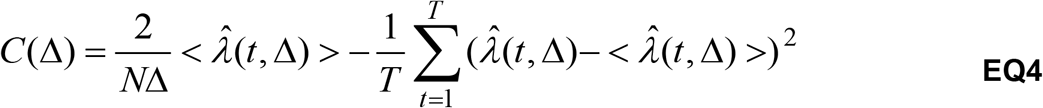

where 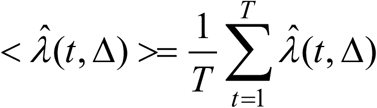

The performance of this cost function will be contrasted with AIC and discussed further in the result section.

### Dependence of optimal bin to the frequency content

The dependence of a histograms optimal bin is theoretically worked out using a similar measure of optimality as MSE (Scott 1979). When adapted to our problem and assuming a continuous time firing rate, the dependency of the optimal binsize to the true firing rate is as following:

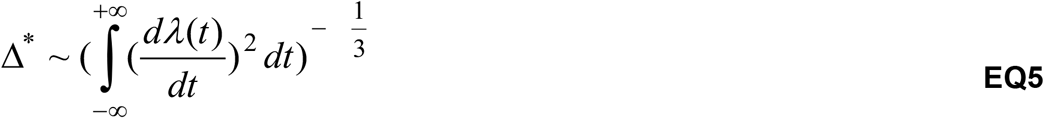

Using Parseval’s theorem and Fourier transform properties we can rewrite **EQ5** in the frequency domain as following:

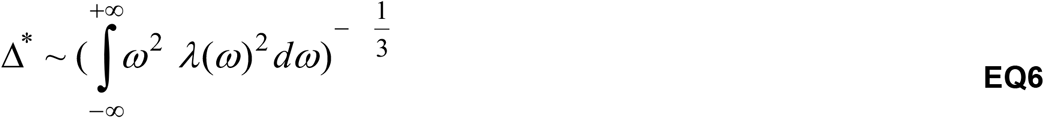

For the sinusoidal firings considered here **EQ6** can be solved explicitly and shows the optimal binsize to be proportional to 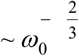 where *ω*_0_ represents the frequency of the sinusoid. This can in fact be verified in **FIG 3.b** where the power law matches the changes of MSE over the range of simulated frequencies. More generally for firing rates that have relatively constant power up-to a cutoff frequency the optimal binsize will be simply proportional to the inverse of the cutoff frequency or alternatively 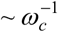 with *ω*_*c*_ being the cutoff.

### Behavior and electrophysiology

Rats were trained with auditory cues (up to 10s) randomly presented on a variable interval schedule with an average interval of 60s. A cartoon of the experimental apparatus is shown in **FIG 5a** (For a detailed description of the experimental paradigm see (Nicola et al. 2004)). Pressing the active lever only during presentation of the cue resulted in the termination of the cue and delivery of a 10% sucrose reward into an adjacent reward receptacle (**FIG 5b**). Another cue (not shown) was interspersed with DS presentation but was not rewarded and is not used for our analysis here. The timing of cue presentations, lever-presses and the entries into and exits out of the reward receptacle were all simultaneously recorded together with the firing of ventral striatal neurons using chronically implanted micro-array electrodes. A sub-set of randomly chosen neurons (having a baseline firing rate >1Hz, n=64) from a previous study is used here (Ambroggi et al. 2011).

### Simulation and data analysis

All simulations and data analyses were done in MATLAB 7.9.0. For all the firing rate simulations, the spike trains are generated using a Poisson point process with a temporally changing rate. The algorithms used for simulation and analysis are available upon request.

## Results

### Optimal Binsize Motivation

It is generally accepted that changing the bin duration can affect the quality of the firing rate estimation when making PETHs (Endres et al. 2010; Shimazaki and Shinomoto 2007; Shinomoto 2010). The trade off is between two desirable qualities, one is high temporal precision which favors smaller binsizes and the other one is low noise levels which favors large binsizes. To demonstrate this trade off let us consider two rather extreme examples, one is a neuron with a constant firing over time and the other is a neuron with an oscillatory firing. For the first example, we assume to have a Poisson point process with a constant firing rate. Although in this case the firing rate is constant, the timing of spikes in each trial will be variable as shown in **FIG 1a**. Therefore one expects that using larger binsizes can help eliminate the local variation in the inter spike intervals while small binsizes overfit the spiking data resulting in noisy estimates of the firing (**FIG 1b**). The MSE between the PETH estimates using different binsizes and the true firing rate shows that in this case the estimation error decreases by increasing the binsizes (**FIG 1c**). Therefore in this example the largest binsize possible is the optimal binsize. However, this conclusion does not hold for the second example neuron (**FIG 1d-f**). Here this neuron changes its firing rate during a trial as shown in **FIG 1d**. Unlike the previous example the choice of excessively large binsizes for this neuron results in loss of temporal resolution in estimating the firing rate for this neuron. On the other hand, choosing a very small binsize also results in noisy estimates by over fitting the local jitters in firing (**FIG 1e**). This is in fact what we observe after plotting the MSE between the estimates and the true firing rate (**FIG 1f**). In this case, the optimal binsize takes an intermediate value in the range of binsizes.

**FIG1.**
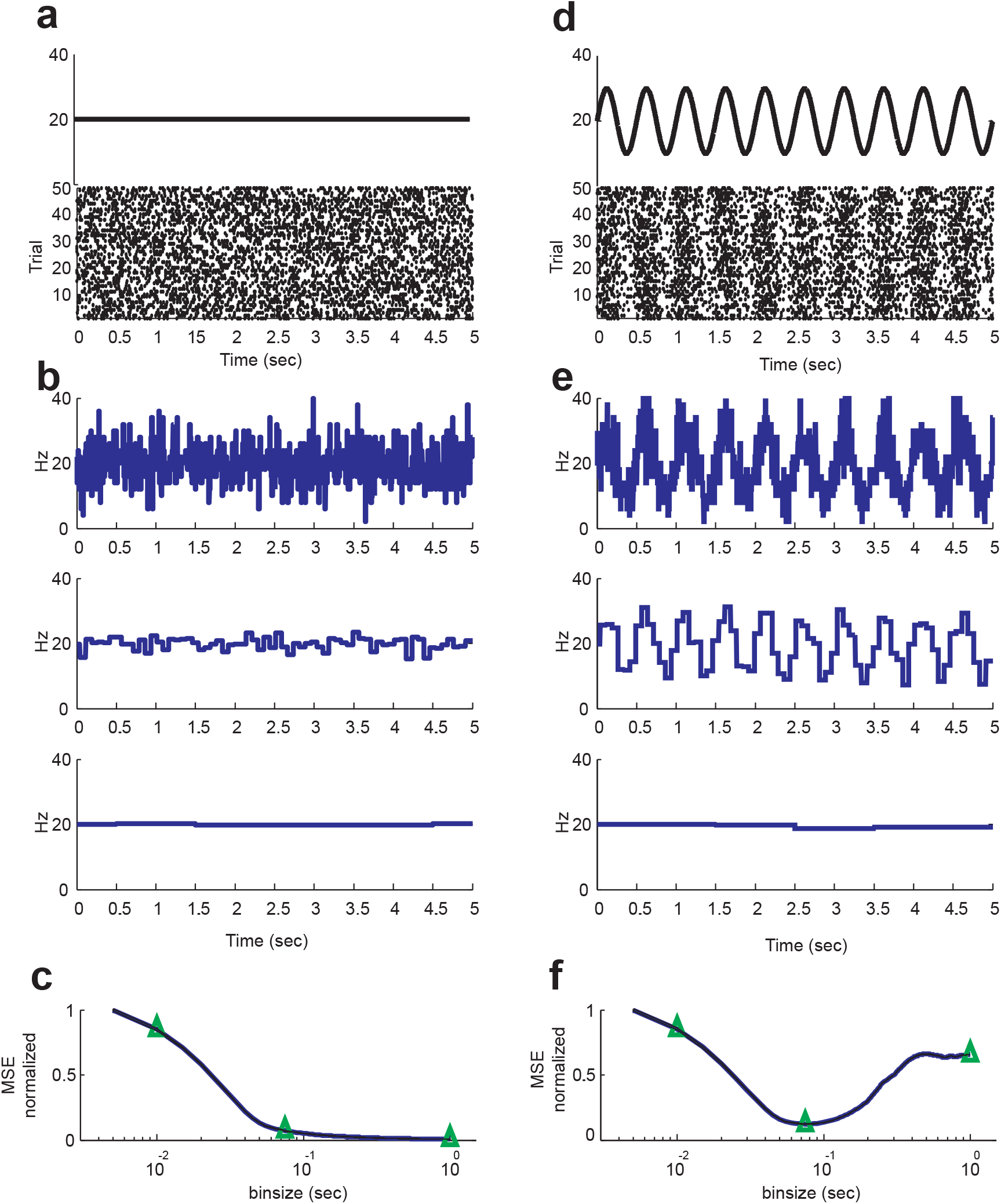
Dependence of Firing Rate Estimate on the Binsize Choice. (**a**) Firing rate and simulated spike train raster with constant Poisson rate of 20Hz. (**b**) Firing rate estimate using 10 ms binsize (1st row) 75 ms (2nd row) and 1 sec (3rd row). (**c**) MSE for the constant Poisson firing monotonically decreases by increasing the binsize. Green triangles mark the 10 ms, 75 ms and 1 sec binsizes used in the estimates shown in part A. (**d**) Firing rate and simulated spike raster with a sinusoidal oscillating firing with an average of 20Hz and oscillation frequency of 2Hz. (**e**) Firing rate estimate using 10 ms binsize (1st row) 75 ms (2nd row) and 1 sec (3rd row). (**f**) MSE for the oscillating Poisson firing reaches a minimum at 75 ms. Green triangles mark the 10 ms, 75 ms and 1 sec binsizes used in the estimates shown in panel **c**.

These results clearly demonstrate the importance of choosing the right binsize for accurate estimation of firing dynamics for neurons. It also shows that the optimal binsize can be different for different neurons and therefore questions the validity of population analysis of data with fixed binsizes without regards to needs of individual PETHs. In the next section we investigate the role of factors such as the number of trials and the temporal dynamics of firing on the optimal binsize using simulated data. We also show in dealing with real data, when the true firing rate is not known, the optimal binsize can be approximated by minimizing Akaike AIC as a cost function that behaves similarly to the MSE (see Methods).

### Simulation Results

Given certain regularity conditions, any signal can be approximated as an integral of sinusoids over a range of frequencies (i.e. frequency decomposition) (Oppenheim et al. 1997). Therefore studying the optimal binsize for sinusoidal firing patterns as basis functions should provide insights that are generalizable to other arbitrary firing patterns.

Let us start with the hypothetical neuron shown in **FIG 2a**. As shown in the figure this neuron has an oscillatory firing with an average of 20 spikes/sec and a sinusoidal modulation of ±10 spikes/sec at 1Hz. Let us further assume that we have recorded the activity of this neuron in 50 trials of 5 second durations each. As can be seen over trials the neurons activity follows the general rise and fall of the underlying sinusoidal rate while the details of spike timings vary from trial to trial as typically seen with real neuronal firing.

**FIG2.**
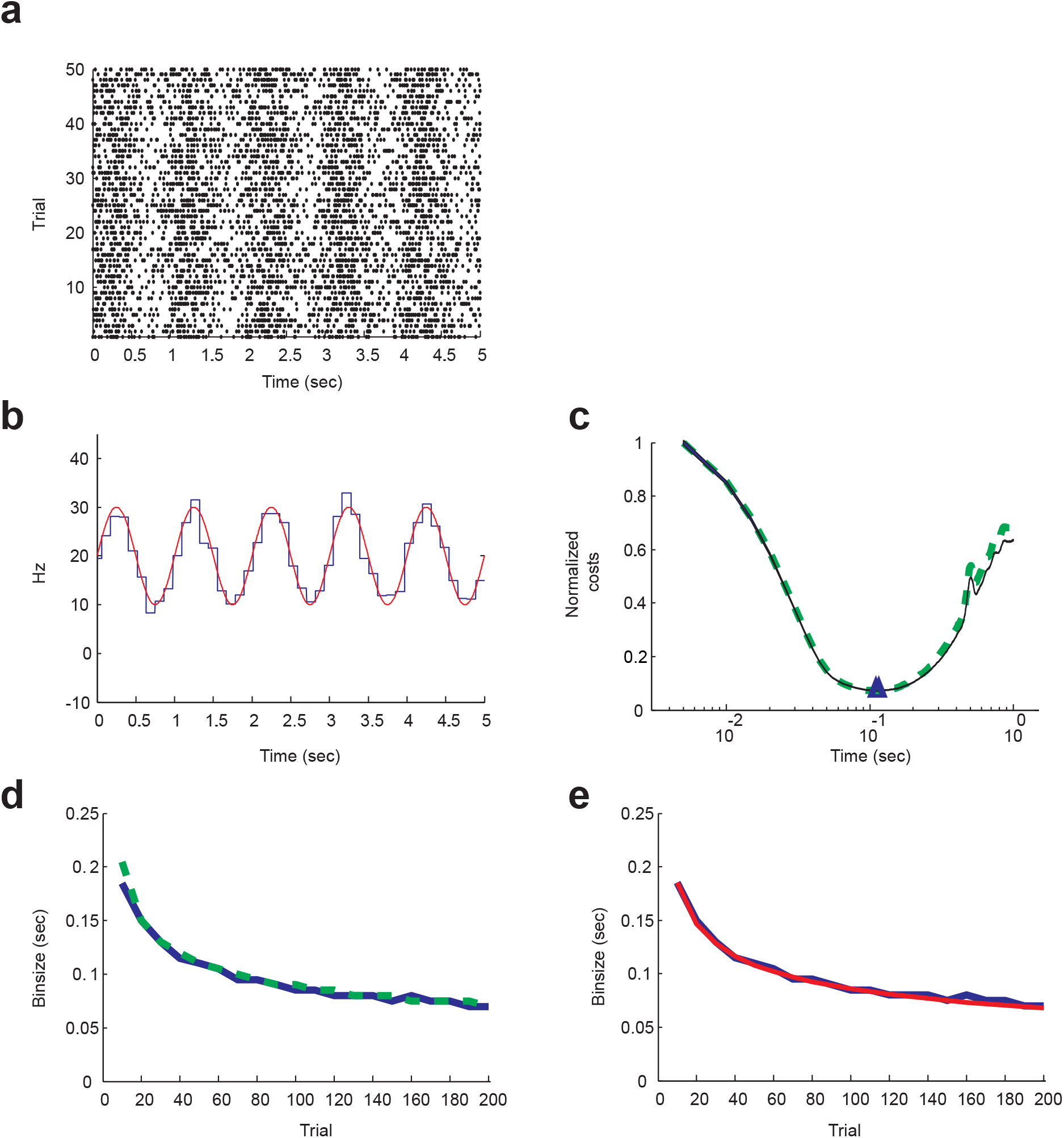
Optimal Binsize Using AIC and Its Dependence on the Number of Trials. (**a**) Simulated spike train raster with a sinusoidal oscillating firing rate with an average of 20Hz and oscillation frequency of 1Hz. (**b**) Firing rate estimate using the optimal binsize (blue) and the true sinusoidal firing rate (red) overlaid. (**c**) MSE (solid blue) and AIC costs (dashed green) as a function of the binsizes overlaid. AIC cost over binsizes closely matches MSE. (**d**) Decrease of the optimal binsize as estimated by MSE (solid blue) and by AIC (dashed green) as a function of increasing trial numbers. Changes of the optimal binsize as predicted by MSE and AIC closely match each other. (**e**) Decrease of the optimal binsize as a function of increasing trial numbers as predicted theoretically (solid red) matches the estimates made by MSE (solid blue).

**FIG3.**
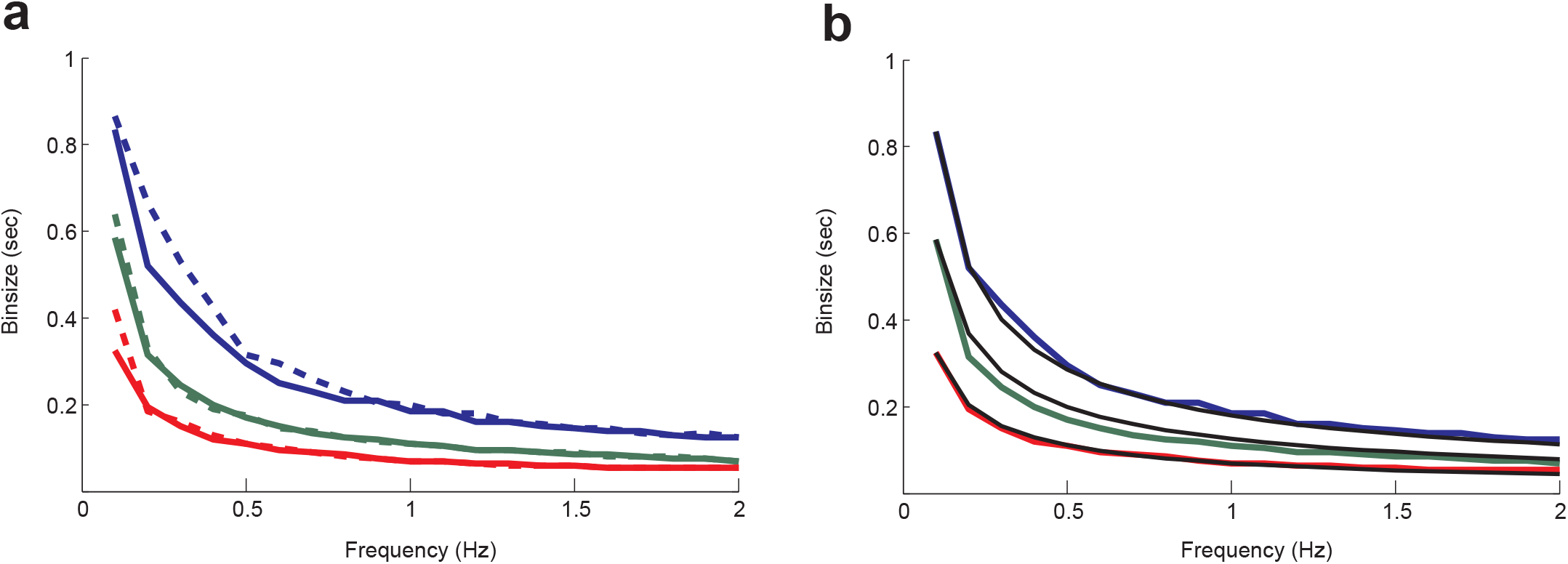
Codependence of the Optimal Binsize on the Firing Rate Bandwidth and the Trial Number. (**a**) The optimal binsize decreases by increasing the frequency of the firing rate changes (0.1 to 2 Hz) and also by increasing the trial numbers (10, 50 and 200 shown by blue, green and red respectively). Solid lines are based on MSE and the dashed lines are based on AIC. (**b**) The decrease of the optimal binsize by increasing either the trial numbers or the firing rate frequency as predicted theoretically (black) matches the estimates made by MSE (colors are the same as panel **a**).

To estimate the underlying firing rate, we can construct firing rate histograms using varying binsizes and calculate their MSE in estimating the true sinusoidal firing rate. As expected too large or too small of a binsize fail to capture the fluctuations of the firing rate over time as shown by the changes of MSE over binsize values in **FIG 2c**. If we now use the spike timings shown in **FIG 2a** to calculate and plot the AIC cost (see **EQ3**) over the same binsize range, one can see that in fact the changes of AIC and MSE as function of binsize are quite similar (**FIG 2c**). The optimal binsize estimate based on AIC also closely matches the estimate base on MSE (110ms and 115ms for AIC and MSE, respectively). This result importantly shows the possibility of accurately predicting the effect of binsize change on firing rate estimation without using or knowing the true firing rate. The PETH made with the optimal binsize is shown in **FIG 2b** as the best approximation of the true sinusoidal firing.

In general the value of the optimal binsize is a function of number of trials available. One expects that increasing the number of trials should decrease the optimal binsize as more trials improve the signal to noise ratio by cancelling the noise over trials thus allowing smaller binsizes to be used. For the example neuron shown in **FIG 2** simulations show that this in fact is the case. The optimal binsize minimizing the MSE decreased from 180ms with only 10 trial to less than 70ms with 200 trials (**FIG 2d** solid blue). Once more we can see that over this range of trial numbers the optimal binsize based on the AIC closely follows the trend of changes in the optimal binsize based on MSE (**FIG 2e** dashed green). In fact, the dependence of optimal binsize of data histograms on the sample size has been studied theoretically in the field of nonparametric density estimations as far back as 1979 (Izenman 1991; Scott 1979). The result of those studies which can be generalized to firing rate histograms mostly suggest that the optimal binsize decreases proportional to 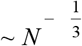 with *N* being the number of trials (or samples in their case). This is slower than the rate of the reduction of noise power which changes as 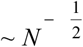 but in fact matches the reduction of optimal binsize seen in the simulation data (**FIG 2e**).

Besides the dependence on the trial number, the optimal binsize should also show dependence on the firing rate changes during a trial. Again the intuition here is that the faster the firing rate changes over time the smaller the optimal binsize should be in order to capture the temporal changes. **FIG 3** shows the dependency of the optimal binsize on the frequency of the sinusoid used in our simulations. The optimal binsize based on MSE and AIC (solid and dashed line respectively) show an inverse relation to the firing rate frequency. Furthermore over the range of frequencies used AIC based optimal binsize is in good agreement with MSE based optimal binsize (**FIG 3a**). The dependence of optimal binsize to frequency of firing rate changes can also be derived theoretically and can be shown to be proportional to 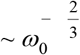 where *ω*_0_ represents the frequency of the sinusoid (see Methods **EQ6**). This can in fact be verified in **FIG 3b** where the power law is shown to match the changes of MSE over the range of simulated frequencies.

### Comparison with other methods

In an elegant study by Shimazaki and Shinomoto, the authors mathematically derived a cost function based on the MSE for firing rates that follow Poisson distribution (Shimazaki and Shinomoto 2007). This cost function adapted to our notation is shown in **EQ4**. However, as they point out, their method diverges for slow fluctuating rates if the number of trials is not large enough. This is in part due to the fact that their derivation is an approximate solution to the problem of minimizing the MSE where the expectation in **EQ1** is replaced by the trial average. Therefore at its current formulation Shimazaki-Shinomoto method should converge to the true MSE given a large number of trials based on the law of large numbers. AIC on the other hand tries to minimize the MSE over all possible observations and as such can provide a superior estimate. As can be seen in **Figure 4** the Shimazaki-Shinomoto cost function tends to overestimate the optimal bin size for slow changing firing rates and low trial numbers. Increasing the number of trials brings this error down for a given frequency (**FIG 4a**). This overestimation of the binsize for slow changing firing rates (lower frequencies) translates into a larger MSE when using the Shimazaki-Shinomoto cost function compared to the AIC although both methods seem to perform equally well for fast changing firing rates (**FIG 4b,c**).

**FIG 4.**
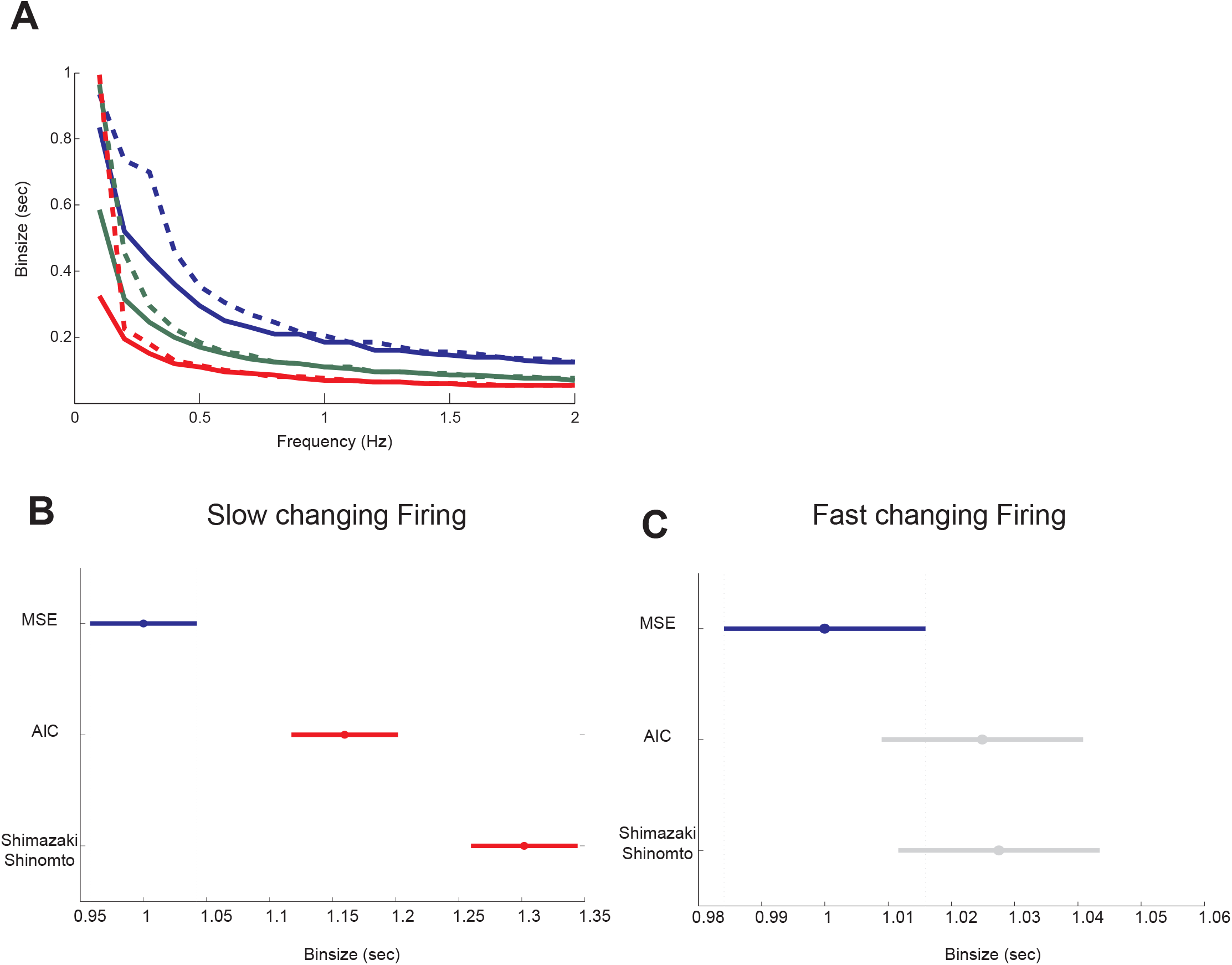
Comparison of AIC with Shimazaki-Shinomoto Method. (**a**) Comparison of the optimal binsizes as estimated by MSE (solid lines) and Shimazaki-Shinomoto method (dashed lines) over a range of firing rate frequencies and trial numbers. Shimazaki-Shinomoto method tends to overestimate the optimal binsize for low frequency (slowly changing) firing rates. (**b**) Comparison of the average sum of squared error between the true firing rate and the estimated firing rate using optimal binsize estimated by minimizing MSE, AIC and Shimazaki-Shinomoto method for slowly changing firing rates <0.5Hz or low bandwidths and (**c**) for fast changing firing rates>0.5Hz or high bandwidths.

**FIG 5.**
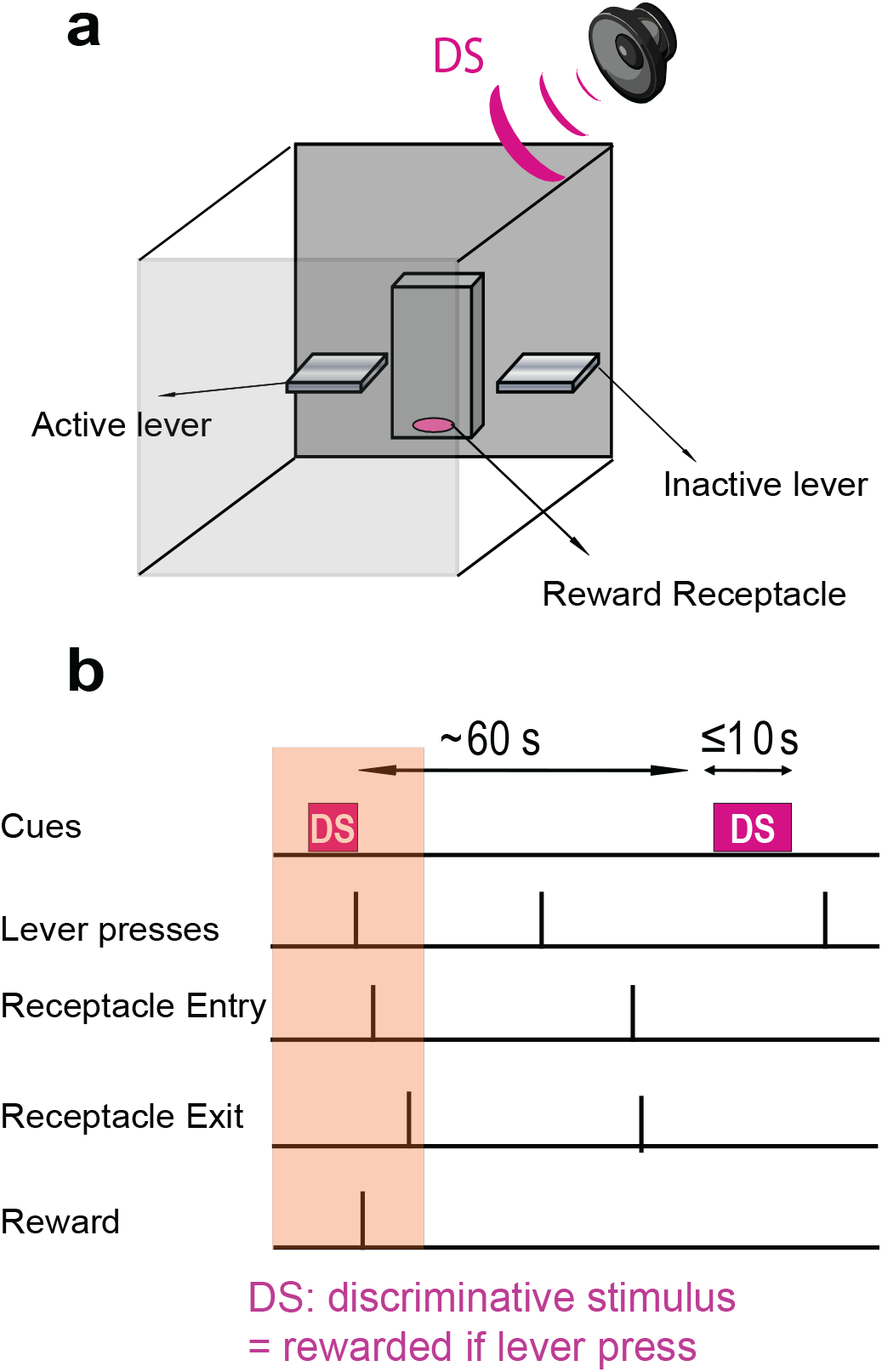
Experimental Procedure. (**a**) Rats were placed in 40 x 40cm chambers containing two levers (active and inactive) on each side of a reward receptacle. A speaker played the discriminative stimulus (DS) above the chamber. (**b**) Behavioral paradigm showing hypothetical sequence of events. Only active lever presses during the DS resulted in reward delivery in the receptacle after which the rat would normally enter the receptacle to consume the reward and exit the receptacle after the consumption. Shaded area illustrates the sequence of rewarded events.

Another study employed an information based technique for optimal firing rate estimation (Paulin and Hoffman 2001). Although they used their method to find the optimal level of smoothing, it can readily be generalized to find an optimal binsize. The measure of optimality used here is closely related to an information theoretic concept known as “cross-entropy” and aims to minimize the disparity between the predicted probability of spiking and the observed probability of spiking in time. This is done by dividing the data into two halves, one used to make predictions and the other for testing. Therefore this approach is equivalent to a cross validation technique using an information based measure. In fact AIC itself is equivalent to a cross validation method (Stone 1974). Although we can verify that their method with some modification should be equivalent to AIC, given the relative complexity of the cross entropy compared to AIC and the computational extensiveness of cross validation, we propose AIC as the simpler and more straightforward measure to be used for finding the optimal binsize.

### Experimental Results

In this section, we demonstrate the effect of varying the binsizes in detection of responses to a reward predictive cue. We used a sub-set of 64 neurons recorded in the nucleus accumbens (NAc) in a discriminative stimulus task (Ambroggi et al. 2011). Briefly, two auditory cues, discriminative stimuli (DS) and neutral stimuli (NS) were randomly presented (with durations of up to 10 s for the DS and 10 s for the NS) on a variable interval schedule with an average interval of 30 s. A lever press was required during DS presentation to cause the delivery of a reward into an adjacent reward receptacle and responding during NS was never rewarded. For simplicity, only the DS is displayed in **Figure 5** with intervals between the two DS’s shown on average to be 60s. (For a detailed description (Nicola et al. 2004))

Responses of neurons to the DS can be obtained by making PETHs time-locked to the cue presentation. Significant changes in the firing rate of the neuron were detected whenever the firing rate of the neuron fell outside the 95% percentile of the baseline (pre-cue) firing rate. Thus excitations are defined as significant increases and inhibitions are defined as significant decreases in the firing rate. **Figure 6a** shows the spike rasters of 5 example neurons that show changes in their spiking rate following the DS presentation. It can be seen that using small binsize of only 10ms for the neurons results in noisy estimates of their firing rate (**FIG 6b**). The noisiness of the estimate seem to especially affect the detection of inhibitory responses that follow the cue (neurons #1,2,3,5). This is because the depression in the firing rate fails to reach the threshold for significance. In some of these cases, the threshold for inhibition is at zero which makes it impossible to detect any significant inhibitions (floor effect for neurons #2,3,5). The noisiness of the estimate not only reduces the detection of inhibitions, thus increasing the false negative error rate, but can also increase detection of spurious activity introducing false positive errors. For example in **Figure 6b**, neurons 1,2,3 and 4 all show seemingly short excitatory or inhibitory activity during the baseline or long after the cue onset. These can be considered as false positives after a cross examination of the firing raster. On the other hand using the optimal binsize for each PETH reveals the inhibitory responses to the cue. It also eliminates many of the falsely detected activities thus reducing the false positive rate (**FIG 6d**).

**FIG 6.**
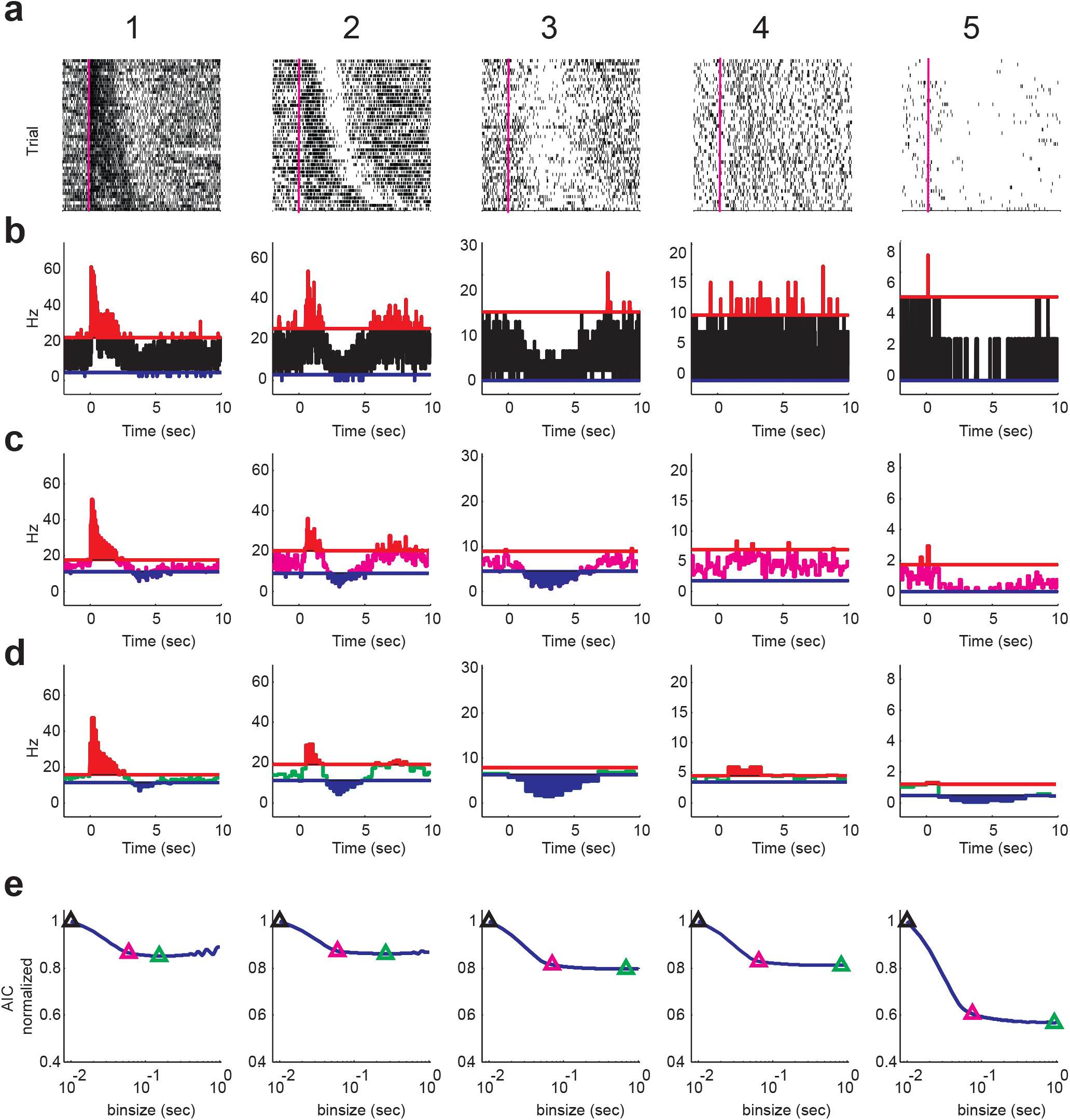
Example PETHs using Small, Deflection Point and Optimal Binsizes. (**a**) Firing raster of example neurons (neurons 1-5 from left to right) and the corresponding PETHs made with (**b**) binsize equal to 10ms, (**c**) binsize equal to the AIC deflection point and (**d**) AIC optimal binsize for each example neuron. (**e**) The AIC cost function for each neuron over a range of binsizes from 10ms to 1 sec. The AIC at 10ms binsize, deflection point and the optimal binsize are marked by black, magenta and green triangles respectively.

Interestingly, the AIC curve shows a rapid decrease for binsizes <~60ms for most neurons shown. This is followed by a deflection point (DP) in the AIC curve after which the changes of the AIC curve over the binsizes become small (**FIG 6e**, magenta and green triangles correspond to DP and optimal binsizes respectively). The optimal binsize which corresponds to the AIC minimum is often much larger than the DP binsize. Although using the optimal binsize results in the best estimate of the firing rate as shown in **FIG 6d** and **FIG 6e**, due to rather large sizes (>500ms for neurons 3,4,5) it may result in insufficient temporal resolution when a more precise detection of changes in the firing is required. Since the AIC gain for using optimal binsize over the DP is rather small (often <10% decrease in AIC cost) in cases where temporal precision is important like detecting the response onset one might choose to use the DP instead of the optimal binsize. However it should be noted that using DP can yield suboptimal results in response detection. For example while it outperforms the minimum binsize in revealing the inhibitions for neurons 1,2 and 3, in the more noisy cases for neurons 4 and 5 it fails to detect the excitation and inhibition to the cue (**FIG 6c**). Cross examination of rasters in **Figure 5a** show the presence of a weak excitation in neuron 4 and an a weak inhibition in neuron 5 following the cue which is correctly detected if optimal binsizes are used (**FIG 6d**).

AIC curves shown in **FIG 6e** also demonstrate that that the optimal binsize is different for each of the neurons shown. Neurons that have a more robust and less noisy response have a smaller optimal binsizes (neurons 1 and 2 with 155ms and 265ms optimal binsizes, respectively) while for neurons with more noisy firings larger optimal binsizes are required to reveal the significant fluctuations in their rate (neurons 3, 4 and 5 with 685ms, 835ms and 940ms optimal bins respectively). On the other hand the DP binsize which turns out to be roughly the same for all neurons in the task (~60 ms) marks a lower bound on the level of temporal resolution that is possible for this population of neurons before the AIC cost gets prohibitively large.

As discussed previously, based on theoretical (**EQ6**) and simulation results (**FIG 2,3**) the optimal binsize for each neuron depends on the number of trials. If the neuronal response is stationary over trials (i.e. its response only depends on the time locked event but not trials) the optimal binsize should be a decreasing function of the trial number following the previously discussed 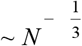 power law. However there are often other behavioral events that happen with variable timing in trials relative to the time locked event that can cause significant non-stationarities. For instance we have shown previously that PETHs like the ones shown in **Figure 6** are often modulated by other temporally close events with variable timing in each trial (Ghazizadeh et al. 2010). Therefore one would predict that the reduction of optimal binsize with increasing trials for real data should be much lower than the theoretical rate. One way to estimate the dependence of optimal binsize for the PETHs of a given neuron is by choosing various random subsets of the available trials and calculate the resulting optimal binsize for a given trial number. Performing this analysis on our experimental data shows that on average, the optimal binsize decreases by adding more trials to the experiment. However, as expected the rate of this decrease is much slower than predicted for stationary rates (**FIG 7a**). The distribution of the decrease rates of optimal binsize over trial numbers for all the neurons also shows that while some neurons follow the theoretical value of 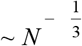 most neurons show a slower decrease as a function of trial number while others show no change or even an slight increase of the optimal binsize as a function of trial number (**FIG 7b**). This results in average rate being ~*N*^−0.1^ for this population of neurons.

**FIG 7.**
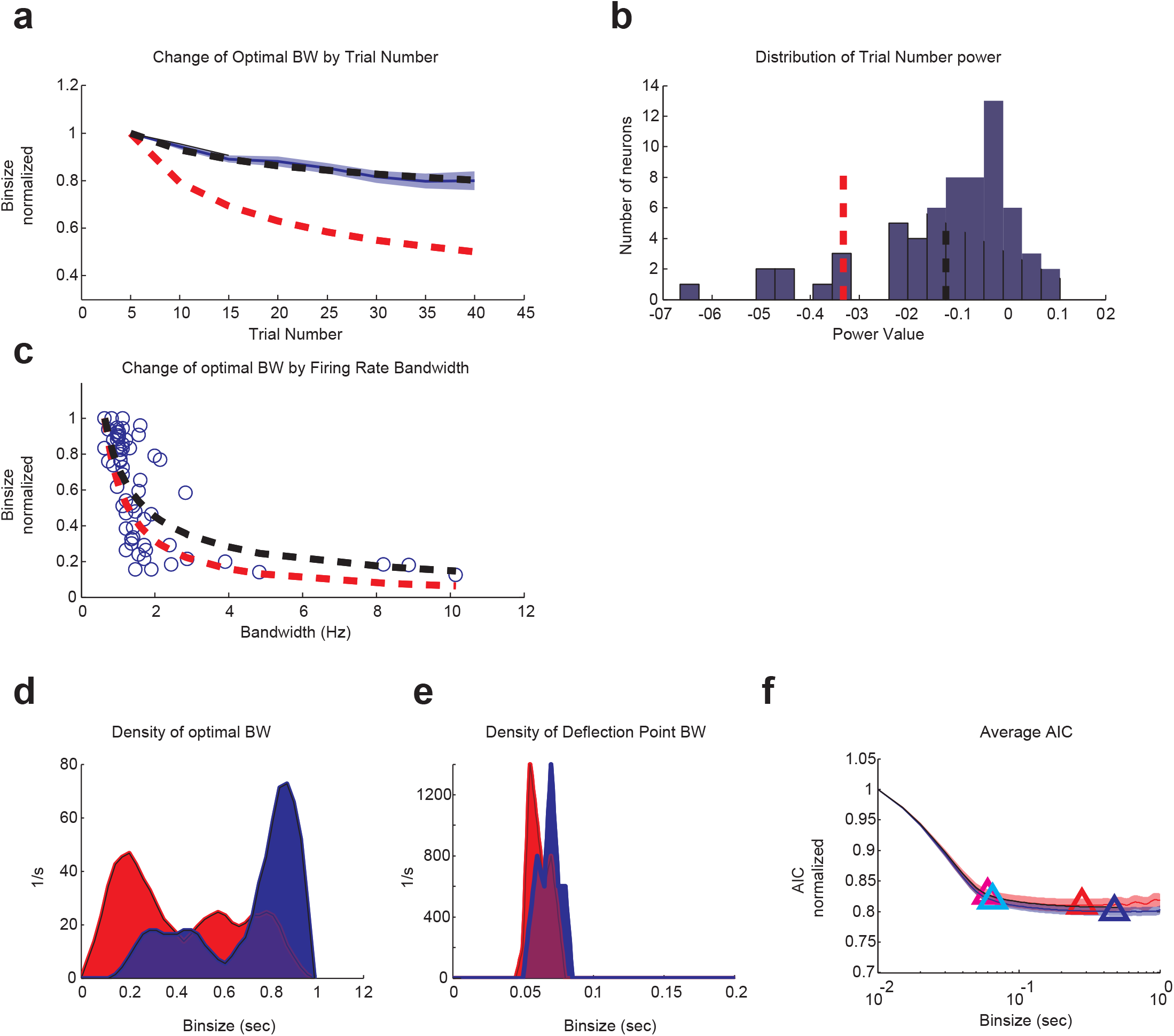
Distribution of Optimal Binsize in the Recorded Neurons and Its Codependence on the Trial Number and Firing Rate Bandwidth. (**a**) The optimal binsize decreases on average for the recorded neurons as the number of trials increases (solid blue: average optimal binsize, blue shade: standard error of the mean). The theoretical rate of decrease (dashed red) and the polynomial fit to the data (dashed black) showing a slower rate of decay for actual data compared to what is predicted for an idealized stationary point process. (**b**) The distribution of the rate of decay of the optimal binsize by the number of trials. Negative values indicate decreasing and positive values indicate increasing with larger trial numbers. Theoretical rate of decay (dashed red) and the average rate from the recorded data (dashed black). (**c**) The optimal binsize decreases on average for the recorded neurons as bandwidth of the firing rate increases. Each open circle corresponds to one neuron. The theoretical rate of decrease (dashed red) and the polynomial fit to the data (dashed black) showing a similar rate of decay for recorded data compared to what is predicted. (**d**) Optimal binsize density for all cue responsive neurons overlaid for excitations (red) and inhibitions (blue) to the cue. (**e**) DP binsize density for all cue responsive neurons overlaid for excitations (red) and inhibitions (blue). (**f**) Average AIC cost for neurons with excitatory responses (red) and inhibitory responses to the cue. Minimum and deflection point of the average AIC for excitation and inhibition is marked with triangles (blue and red: optimal for excitation and inhibition respectively, cyan and magenta: DP for excitation and inhibition respectively)

Another factor that is known to affect the optimal binsize is the bandwidth of the firing rate. Firing rates that have larger and more frequent changes require smaller binsizes to capture their temporal variation compared to firing rates with slower changes. Although we do not have access to the true firing rate of our neurons we should be able to get a relatively accurate estimate of their bandwidth by using the firing rate estimates made with the optimal binsize. Plotting the optimal binsize of each neuron as a function of its firing bandwidth in fact shows a relatively good agreement with the theoretical values as the optimal binsizes are shown to be a decreasing function of the bandwidth with a rate proportional to 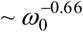 is close to theoretical rate of 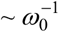 (**FIG 7c**).

The optimal binsize shows a bimodal distribution over our neuronal population with two peaks, one around 0.2sec and another around 0.9sec. Further separation of the binsize distribution between the neurons excited to the cue and inhibited to the cue shows that in fact the peak with the smaller binsize corresponds mostly to excitatory responses while the second peak with the larger binsizes corresponds mostly to inhibitory responses (**FIG 7d**). The distribution of the DP binsize on the other hand shows a much narrower spread around 50ms for excitations and 70ms for inhibitions (**FIG 7e**). The average AIC cost for excitatory and inhibitory responses also demonstrates that the optimal binsize of the average AIC for inhibitions are larger than for excitation with the difference diminishing for the DP binsize (**FIG 7f**).

As discussed previously for the examples in **Figure 6**, the choice of binsize can greatly affect the detection of responses and their temporal dynamics including the magnitude, onset and offset of excitations and inhibitions. To further examine this effect, we detected response to DS using both the optimal binsize and a fixed 60ms (DP-like) binsize. Since the value of DP binsize was very similar for all neurons and was shown to be around 60ms (**FIG 7e**) the choice of this binsize should give similar results to DP binning while being the same for all neurons. Even though similar number of responsive neurons is detected using 60ms or optimal binsize (49 and 54 for OP and 60ms respectively) the profiles of detected cue responses look different in each condition (**FIG 8a**). Using the optimal binsize generally results in much higher z-scores in the responsive neurons compared to the baseline mainly due to reduction of baseline noise. This is true for both excitatory and inhibitory responses. The average z-score for each response type shows a marked increase by using the optimal binsize over 60ms binsize (**FIG 8b**). Inhibitory responses are specifically enhanced in their z-score after using the optimal binsize resulting in a more symmetrical look in the amplitude of excitatory and inhibitory z-scores using optimal binsizes for each neuron.

**FIG 8.**
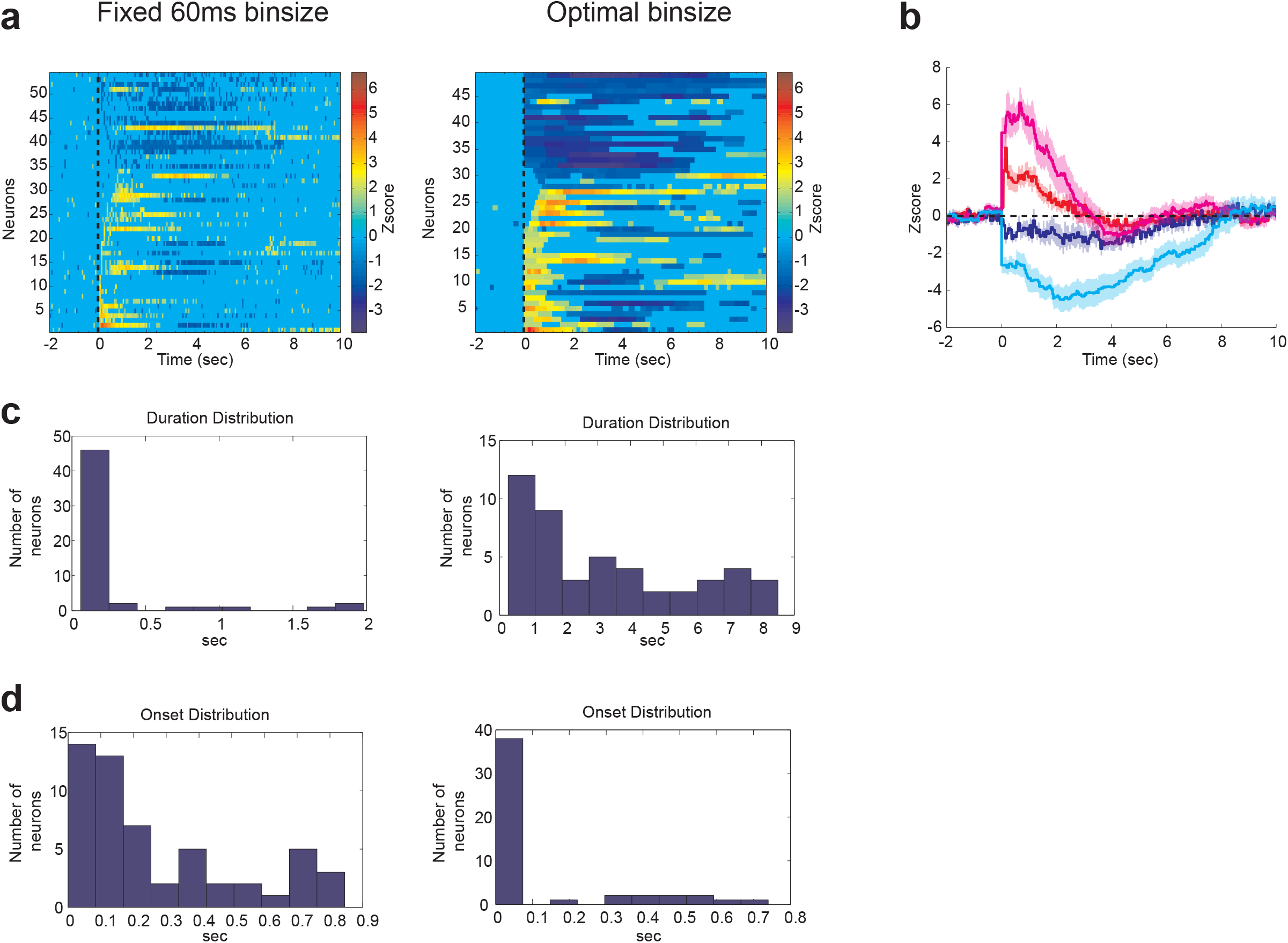
Comparison of Population Profile of Responses to the Cue Based on PETHs Made with Optimal Versus fixed 60ms Binsizes. (**a**) Color coded PETH z-scores made with fixed 60ms (left column) and with optimal binsize (different for each neuron) (right column) for all the responsive neurons to the cue. Each line in the color coded plots corresponds to the firing of one neuron in z-score. Only z-scores outside the 90% confidence interval of baseline (pre-cue firing) are color coded. Values within the confidence interval are all colored at zero. (**b**) Average PETH z-scores made with fixed 60ms binsize versus z-scores made with optimal binsize. Red and blue correspond to excitatory and inhibitory responses made with 60ms binsize respectively. Magenta and cyan correspond to excitatory and inhibitory responses made with optimal binsize respectively. (**c**) Distribution of the cue response durations made with fixed 60ms (left column) and with optimal binsize (right column) for all the responsive neurons to the cue. (**d**) Distribution of the cue response onsets made with fixed 60ms (left column) and with optimal binsize (right column) for all the responsive neurons to the cue.

The distribution of response durations also shows that using the optimal binsizes results in detecting an array of durations from less than 1 sec to more than 8 sec while most of the response durations detected by 60ms binsize are less than 0.5 sec (**FIG 8c**). This is due the discontinuities observed in the profile of responses using 60ms binsize for each neuron as shown in **FIG 8a**. On the other hand due to the relatively large values of the optimal binsizes the distribution of the response onsets using optimal binsizes lacks the resolution to show the natural variation seen when using smaller binsizes. Therefore using a small DP like binsize to make PETHs has a higher resolution for the onset detection and results in more accurate estimates of the response onset for this dataset (**FIG 8d**).

## Discussion

The main goal of systems neuroscience is to relate the firing of neurons in the brain to behavioral or environmental events. PETHs are widely used in the field for exactly this purpose and have contributed in a major way to our understanding of the nervous system. The usefulness of PETH depends on at least two qualities. First is how well the PETH manages to cancel the firing variability across time and trials to reveal the firing rate modulations related to the event of interest. Second is how well it can detect transient changes in the firing rate that are not due to noise. Maximizing both of these functions is often not possible since noise cancellation often requires use of large binsizes for a PETH that in turn reduces the temporal resolution (**Fig 1**).

We have addressed this trade off between noise and temporal resolution by proposing a principled way for choosing the optimal binsize for each PETH by using the AIC method. Briefly this method defines a cost with one term that favors smaller binsizes and another term that favors larger binsizes to reduce the total number of bins (taken as the number of model parameters). Simulation results show that PETHs made with the AIC optimal binsize often have the lowest MSE errors when compared to the true firing rates.

A PETH optimal binsize depends on at least two factors, one that is intrinsic to the neural activity being estimated and the other that is controlled by the experimenter. The first factor is how much and how fast the firing of a neuron shows changes to the event of interest. The faster these changes happen in the firing the smaller the binsize needs to be to capture the temporal dynamics. Theoretical results show that the optimal binsize is almost inversely correlated with the frequency content of the firing rate modulation. We find good agreement with theory in simulated (**FIG 3b**) as well as actual data (**FIG 7c**). The second factor affecting the optimal binsize is the number of trials used to make a PETH. Intuitively the higher the number of trials is the smaller the optimal binsize can be. This is because a good degree of noise cancellation can happen over trials rather than time. Theoretical results estimate the relation between trial number and optimal binsize to follow an inverse cube power law. Our simulation results are again in good agreement with this result (**FIG 2e**). An important assumption for this relationship is that the true neuronal firing to the event of interest should not change from trial to trial (stationarity). However, the mere presence of other variably occurring events that can modulate the firing of the neuron during a trial nullifies this assumption when dealing with actual neural data (Ghazizadeh et al. 2010). In line with the violation of stationarity, we find that the actual decrease rate of the optimal binsize as a function of trial number is often much slower than inverse cube power (−0.33) averaging to about −0.1 in our data sample (**FIG 7a,b**).

We also show that the binsize choice has a dramatic impact on response detection. Many false positives or false negatives can result if a binsize that is too small or too large is used. As stated before, the optimal binsize for each PETH differs according to the number of trials and firing dynamics. This difference in optimal binsizes means that using the same binsize for analysis of neural populations can lead to misclassification or misinterpretation of the data. In our population of neurons in the NAc which are considered to be mainly low firing medium spiny neurons (Berke et al. 2004; Mallet et al. 2005), this misinterpretation can particularly occur when comparing excitations and inhibitions. Due to the floor effect of low baseline firing and the slower dynamics of inhibitions, inhibitory responses are only revealed at larger binsizes (>0.5). In fact using binsizes that are optimal for detection of excitations often result in missing weak inhibitions in the neuronal population (**FIG 6**). Analysis of optimal binsize across excitations and inhibitions clearly show this separation of optimal binsizes across these two response groups (**FIG 7d-f**). Interestingly the use of optimal binsize for both excitations and inhibitions showed that the average z-score magnitude of inhibitions can be comparable to excitations in the NAc, a fact that is missed if equal binsizes are used for all neurons (**FIG 8a,b**).

One draw back using optimal binsizes can be the loss of resolution in estimating the response onset since optimal binsizes are often large (>0.2). One solution is to use the binsize at the deflection point (DP) of AIC cost. This DP binsize happens after the sharp drop of the AIC cost but is usually much smaller than the optimal binsize (**FIG 6e**). We demonstrated that the use of DP binsize can improve the onset estimation while having little effect on the percentage of detected neurons (**FIG 8a,d**). However, for the response duration optimal binsize provides more accurate results compared to short DP binsizes (**FIG 8a,c**).

Recently various methods for making optimal PETHs have been developed and contrasted (Endres et al. 2008; Shinomoto 2010). However many of these methods are based on maximum likelihood parameterizations or cross validations that are computationally intensive (Endres et al. 2010; Paulin and Hoffman 2001; Ritov et al. 2002). In contrast, our AIC method provides a simple recipe for binsize selection and has near optimal results over a wide range of trial numbers and firing rates. Another alternative to AIC is proposed by Shimazaki and Shinomoto and provides a cost function that minimizes the MSE in case of a variable Poisson firing rate (Shimazaki and Shinomoto 2007). Our comparisons show that AIC outperforms this method especially in the case of slow changing firing rates and low number of trials (**FIG 4**).

In conclusion, we demonstrated the importance of the binsize choice in accuracy of firing rate estimations using PETHs. We proposed that optimal binsize for a PETH can be estimated using an AIC cost and we investigated the dependence of this optimal binsize on other factors such as firing rate modulation and trial number. Furthermore our results show that for neurons with low basal firing rate like NAc medium spiny neurons small binsizes can miss many of the inhibitory responses to the events. Accordingly optimal binsizes for inhibitions turn out to be much larger than excitations. Therefore we propose use of optimal binsize for PETH based response detection.

## Acknowledgments

This project has received funding from the ATIP/Avenir program, the Wheeler Center for Neurobiology of Addiction to FA.

We thank Gregory Hjelmstad for critical commenting on the manuscript and Howard L. Fields for his incommensurable support and mentoring.

